## Supplementary information: this files contains the supplementary figures and associated figure legend. for "An atypical F-type ATPase is necessary for the function of the antibody cleavage system MIB-MIP in mycoplasmas"

**Supplementary Figures Legends:**

Fig S1: Structure predictions of the F_1_-like X_0_ ATPase sub-units. The 3D structures of the different proteins encoded in the F_1_-like X_0_ ATPase locus have been predicted using the AlphaFold3 model. The amino-acid sequence corresponding to the mnemonic MMCAP2_0581-MMCAP2_0575 was used as input. The first model generated for each prediction is presented. The name of each protein is based on the last four characters of the corresponding locus mnemonics. The 3D structures were visualized using ChimeraX and are shown as coloured ribbons (colouring is according to the locus map presented in Figure 1B). When possible, structural comparison to the F_1_F_0_ ATPase sub-units of *E. coli* (PDB: 6OQR) where performed using ChimeraX and the *matchmaker* command. Amino-terminal and Carboxyl-terminal positions are indicated as Nt and Ct, respectively. **A**) Predicted structures of the F_1_-like sub-unit monomers (top) and comparison to their cognate F_1_ sub-units (bottom, in grey). **B**) Predicted structure of the F_1_-like complex generated using AlphaFold3 multimer by inputting multiple copies of several sequences (1x 0579; 1x 0578; 3x 0576; 3x 0575). The predicted complex is presented in two orientations rotated by 90° (left: lateral view; right: axial view). **C**) Predicted structures of the X0 sub-unit monomers. **D**) Predicted structure of the F_1_-like X_0_ complex generated using AlphaFold3 multimer by inputting multiple copies of several sequences (1x 0581; 3x 0580; 1x 0577; 1x 0579; 1x 0578; 3x 0576; 3x 0575). The predicted complex structure (left) is presented alongside the experimentally acquired structure of the F_1_F_0_ ATPase of *E. coli* (right, grey, PDB: 6OQR). Scalebar: 10 Å.

Fig S2: Phylogenetic analysis of the predicted homologs of MMCAP2_0581. Phylogenetic analysis of the predicted homologs of MMCAP2_0581 was performed using the Phylogeny tool (<https://www.phylogeny.fr/>) in “One click” mode. The output phylogenic tree was visualized using iTol (https://itol.embl.de/) as a circular tree. The position of MMCAP2_0581 in the tree is marked by the coloured arrowhead. *Mycoides* cluster and the *Bovis*-*agalactiae* cluster are highlighted in blue, showing their phylogenetic proximity, in accordance with a horizontal gene transfer scenario, and similarly for the *Hominis* and *Ureaplasma* clusters that are highlighted in green.

Fig S3: Phylogenetic analysis of the predicted homologs of MMCAP2_0580. Phylogenetic analysis of the predicted homologs of MMCAP2_0580 was performed using the Phylogeny tool (<https://www.phylogeny.fr/>) in “One click” mode. The output phylogenic tree was visualized using iTol (https://itol.embl.de/) as a circular tree. The position of MMCAP2_0580 in the tree is marked by the coloured arrowhead. *Mycoides* cluster and the *Bovis*-*agalactiae* cluster are highlighted in blue, showing their phylogenetic proximity, in accordance with a horizontal gene transfer scenario, and similarly for the *Hominis* and *Ureaplasma* clusters that are highlighted in green.

Fig S4: Phylogenetic analysis of the predicted homologs of MMCAP2_0579 (γ–like subunit). Phylogenetic analysis of the predicted homologs of MMCAP2_0579 was performed using the Phylogeny tool (<https://www.phylogeny.fr/>) in “One click” mode. The output phylogenic tree was visualized using iTol (https://itol.embl.de/) as a circular tree. The position of MMCAP2_0579 in the tree is marked by the coloured arrowhead. *Mycoides* cluster and the *Bovis*-*agalactiae* cluster are highlighted in blue, showing their phylogenetic proximity, in accordance with a horizontal gene transfer scenario, and similarly for the *Hominis* and *Ureaplasma* clusters that are highlighted in green.

Fig S5: Phylogenetic analysis of the predicted homologs of MMCAP2_0578 (ε-like subunit). Phylogenetic analysis of the predicted homologs of MMCAP2_0578 was performed using the Phylogeny tool (<https://www.phylogeny.fr/>) in “One click” mode. The output phylogenic tree was visualized using iTol (https://itol.embl.de/) as a circular tree. The position of MMCAP2_0578 in the tree is marked by the coloured arrowhead. *Mycoides* cluster and the *Bovis*-*agalactiae* cluster are highlighted in blue, showing their phylogenetic proximity, in accordance with a horizontal gene transfer scenario, and similarly for the *Hominis* and *Ureaplasma* clusters that are highlighted in green.

Fig S6: Phylogenetic analysis of the predicted homologs of MMCAP2_0577. Phylogenetic analysis of the predicted homologs of MMCAP2_0577 was performed using the Phylogeny tool (<https://www.phylogeny.fr/>) in “One click” mode. The output phylogenic tree was visualized using iTol (https://itol.embl.de/) as a circular tree. The position of MMCAP2_0577 in the tree is marked by the coloured arrowhead. *Mycoides* cluster and the *Bovis*-*agalactiae* cluster are highlighted in blue, showing their phylogenetic proximity, in accordance with a horizontal gene transfer scenario, and similarly for the *Hominis* and *Ureaplasma* clusters that are highlighted in green.

Fig S7: Phylogenetic analysis of the predicted homologs of MMCAP2_0576 (α-like subunit). Phylogenetic analysis of the predicted homologs of MMCAP2_0576 was performed using the Phylogeny tool (<https://www.phylogeny.fr/>) in “One click” mode. The output phylogenic tree was visualized using iTol (https://itol.embl.de/) as a circular tree. The position of MMCAP2_0576 in the tree is marked by the coloured arrowhead. *Mycoides* cluster and the *Bovis*-*agalactiae* cluster are highlighted in blue, showing their phylogenetic proximity, in accordance with a horizontal gene transfer scenario, and similarly for the *Hominis* and *Ureaplasma* clusters that are highlighted in green.

Fig S8: Phylogenetic analysis of the predicted homologs of MMCAP2_0575 (β-like subunit). Phylogenetic analysis of the predicted homologs of MMCAP2_0575 was performed using the Phylogeny tool (<https://www.phylogeny.fr/>) in “One click” mode. The output phylogenic tree was visualized using iTol (https://itol.embl.de/) as a circular tree. The position of MMCAP2_0575 in the tree is marked by the coloured arrowhead. *Mycoides* cluster and the *Bovis*-*agalactiae* cluster are highlighted in blue, showing their phylogenetic proximity, in accordance with a horizontal gene transfer scenario, and similarly for the *Hominis* and *Ureaplasma* clusters that are highlighted in green.

Fig S9: Conservation analysis of the α-like sub-unit of the F_1_-like X_0_ ATPase. The amino-acid sequences of 51 representative α-like homologs were extracted from Table S2, as well as the sequence of the protein AtpA from *E. coli*, and aligned using ClustalW. The alignment quality scores by residue are plotted in purple, and the Walker A motif is highlighted in red (top). The region of the alignment in the black box is detailed below (bottom). The alignment file was opened using JalView (<https://www.jalview.org/>), and coloured according to the *clustal* palette. The Walker A consensus motif GXXXGKT is noted in grey, aligned with the predicted Walker A motif in the α and α-like subunits. The sequence logo of the Walker A motif was generated using Weblogo (<https://weblogo.berkeley.edu>).

Fig S10: Conservation analysis of the β-like sub-unit of the F_1_-like X_0_ ATPase. The amino-acid sequences of 51 representative β-like homologs were extracted from Table S2, as well as the sequence of the protein AtpD from *E. coli*, and aligned using ClustalW. The alignment quality scores by residue are plotted in purple, and the Walker A motif is highlighted in red (top). The region of the alignment in the black box is detailed below (bottom). The alignment file was opened using JalView (<https://www.jalview.org/>), and coloured according to the *clustal* palette. The Walker A consensus motif GXXXGKT is noted in grey, aligned with the predicted Walker A motif in the β and β-like sub-units. The sequence logo of the Walker A motif was generated using Weblogo (https://weblogo.berkeley.edu).

Fig S11: Generation of the mutant strain *Mmc* ΔATPase. **A**) The locus encoding the MIB-MIP-F_1_-like X_0_ ATPase in *Mmc* 1.1 is presented using the same pattern as in Figure 1 (top). The position of the sequence targeted by the guide RNA pgRNA_0577 is denoted by a black diamond. The final locus in the mutant *Mmc* ΔATPase is also displayed (bottom). A detailed view of the recombination arms used to perform the editing in-yeast, following CRISPR-Cas9 stimulated Homologous Recombination is shown (inset). A recombination patch comprised of 2x45 bp homologous arms was generated from two oligonucleotides. The left arm of the patch corresponds to the TAA stop codon of MMCAP2_0582 and the subsequent intergenic spacer. The right arm of the patch corresponds to the terminator of the putative MIB-MIP-F_1_-like X_0_ ATPase operon. The double recombination is shown by grey parallelograms. **B**) Simplex PCR screening of the yeast transformants. The properly edited locus should yield a 555 bp amplicon. “1.1”: *Mmc* 1.1 gDNA template; “+”: positive control; “-”: no DNA control. **C**) Multiplex PCR screening of the yeast transformants. The complete genome of *Mmc* mutants should yield the same 11 amplicons as in the positive control. “1.1”: *Mmc* 1.1 gDNA template; “WT”: *Mmc* GM12 gDNA template; “-”: no DNA control. **D**) PFGE analysis of the bacterial chromosome carried in yeast after restriction by XhoI. The complete genome of *Mmc* mutants should yield the same 3 large fragments (590, 269 and 226 kpb) as the positive control. “1.1”: *Mmc* 1.1 gDNA. **E**) Simplex PCR screening of the bacterial transplants. The properly edited locus should yield a 555 bp amplicon. The lane 1.1 corresponds to the transplant 1 obtained from the yeast clone 1. The lanes 36.1, 36.2 and 36.3 correspond to the 3 transplants obtained from the yeast clone 36. “WT”: *Mmc* GM12 gDNA template; “+”: positive control; “-”: no DNA control. **F**) Multiplex PCR screening of the bacterial transplants. The complete genome of *Mmc* mutants should yield the same 11 amplicons as in the positive control. “WT”: *Mmc* GM12 gDNA template; “-”: no DNA control. *Note: only the sample lanes relevant to this publication are annotated. The other samples correspond to other Mmc mutants that are not presented in this publication.*

Fig S12: Generation of the mutant strain *Mmc* 0575-recode and *Mmc* 0575-K152A. **A**) The locus encoding the MIB-MIP-F_1_-like X_0_ ATPase in *Mmc* 1.1 is presented using the same pattern as in Figure 1 (top). The position of the sequence targeted by the guide RNA pgRNA_0575 is denoted by a black diamond. The edited locus in the mutants *Mmc* 0575-recode or *Mmc* 0575-K152A is also displayed (middle and bottom). A detailed view of the recombination arms used to perform the editing in-yeast, following CRISPR-Cas9 stimulated Homologous Recombination is shown (inset). A recombination patch comprised of the complete and edited locus MMCAP2_0576-MMCAP2_0576 was generated by PCR. The left arm of the patch corresponds to the complete wild-type locus MMCAP2_0576. The right arm of the patch corresponds to theedited locus MMCAP2_0575-recode or MMCAP2_0575-K152A. The double recombination is shown by grey parallelograms. **B**) Simplex PCR screening of the yeast transformants. The properly edited locus should yield a 2927 bp amplicon. “+”: positive control; “-”: no DNA control. **C**) XbaI restriction screening of the Simplex PCR amplicons. The properly edited locus should yield two bands at 1554 bp and 1373 bp. “+”: XbaI restriction positive control; “-”: XbaI restriction negative control. **D**) Multiplex PCR screening of the yeast transformants. The complete genome of *Mmc* mutants should yield the same 11 amplicons as in the positive control. “1.1”: *Mmc* 1.1 gDNA template; “-”: no DNA control. **E**) PFGE analysis of the bacterial chromosome carried in yeast after restriction by XhoI. The complete genome of *Mmc* mutants should yield the same 3 large fragments (590, 269 and 226 kpb) as the positive control. “1.1”: *Mmc* 1.1 gDNA. **F**) Simplex PCR screening of the bacterial transplants. The properly edited locus should yield a 2927 bp amplicon. The lane 1.1, 1.2 and 1.3 corresponds to the transplants 1, 2 and 3 obtained from the yeast clone 1. “+”: positive control; “-”: no DNA control. **G**) Multiplex PCR screening of the bacterial transplants. The complete genome of *Mmc* mutants should yield the same 11 amplicons as in the positive control. “WT”: *Mmc* GM12 gDNA template; “-”: no DNA control. *Note: only the sample lanes relevant to this publication are annotated. The other samples correspond to other Mmc mutants that are not presented in this publication.*

Fig S13: Serum agglutination assays and Immunoglobulin cleavage assays with the mutant *Mmc* strains. **A**) Agglutination of cells by immune goat serum. The cells were grown in axenic conditions, in absence (top) or presence (bottom) of 2% of serum from a goat experimentally immunized, collected 12 days post-inoculation. The bottom of the micro-centrifuge tubes was photographed (top). After resuspension of the flocculates by inversion of the tubes, a sample was mounted between a glass slide and a coverslip and was imaged on a dark-field microscope (bottom - scale bar: 10 µm. **B**) Western blot analysis of the immunoglobulin Heavy Chain cleavage by the MIB-MIP system. Samples corresponding to either culture supernatant or cell pellet derived from the agglutination assays were separated by SDS-PAGE, then analyzed by Western Blotting using primary antibodies targeting either the goat IgG Fc or the goat IgM Heavy Chain. Intact Heavy Chain and MIB-MIP cleaved Heavy Chain are highlighted by a gray and white arrowhead, respectively.

Fig S14: Generation of the mutant strain *Mmc* Rational Operon 0577-ALFA^96^. **A**) The locus encoding the MIB-MIP-F_1_-like X_0_ ATPase in *Mmc* 1.1 is presented using the same pattern as in Figure 1 (top). The locus in the mutant *Mmc* Rational Operon, generated in a previous study, is also shown (middle). The position of the sequence targeted by the guide RNA pgRNA_0577 is denoted by a black diamond. The final locus in the mutant *Mmc* Rational Operon 0577-ALFA^96^ is also displayed (bottom). A detailed view of the recombination arms used to perform the edition in-yeast, following CRISPR-Cas9 stimulated Homologous Recombination is shown (inset). A recombination patch comprised of two ~290 bp homologous arms was generated by PCR. The left arm of the patch corresponds to 288 first coding bases of MMCAP2_0577. The right arm of the patch corresponds to the next 298 bp of MMCAP2_0577. The two arms are separated by 45 bp encoding the ALFA tag (13x3 bp) flanked on each side by two serine encoding codons (2x3 bp). The double recombination is shown by grey parallelograms. **B**) Simplex PCR screening of the yeast transformants. The properly edited locus should yield a 1012 bp amplicon. “+”: positive control; “-”: no DNA control. **C**) Multiplex PCR screening of the yeast transformants. The complete genome of *Mmc* mutants should yield the same 11 amplicons as in the positive control. “1.1”: *Mmc* 1.1 gDNA template; “-”: no DNA control. **D**) PFGE analysis of the bacterial chromosome carried in yeast. The complete genome of *Mmc* mutants should yield the same 1 large fragment (994 kpb) as the positive control. “1.1”: *Mmc* 1.1 gDNA. **E**) Anti-ALFA tag Western blot screening of the bacterial transplants. For each analyzed transplant, whole cell extracts were generated and separated by SDS-PAGE. Anti-ALFA Western blotting was then performed. The *Mmc* mutants should display a single band at ~85 kDa. The lanes 10.1, 10.2, 10.3 and 12.1,12.2, 12.3 corresponds to the 3 transplants obtained from the yeast clone 10 and 12, respectively. “+” recombinant ALFA-tagged protein; “1.1”: *Mmc* 1.1 whole cell extract. *Note: only the sample lanes relevant to this publication are annotated. The other samples correspond to other Mmc mutants that are not presented in this publication.*

Fig S15: Uncropped original images. The figures of this publication contain images that have been cropped and adjusted for contrast or levels. Cropping has been performed to improve legibility or to remove internal control lanes. Adjustments have been applied homogeneously to all images. Image processing does not modify the interpretation of the images. Original uncropped and unadjusted images are provided for reference.

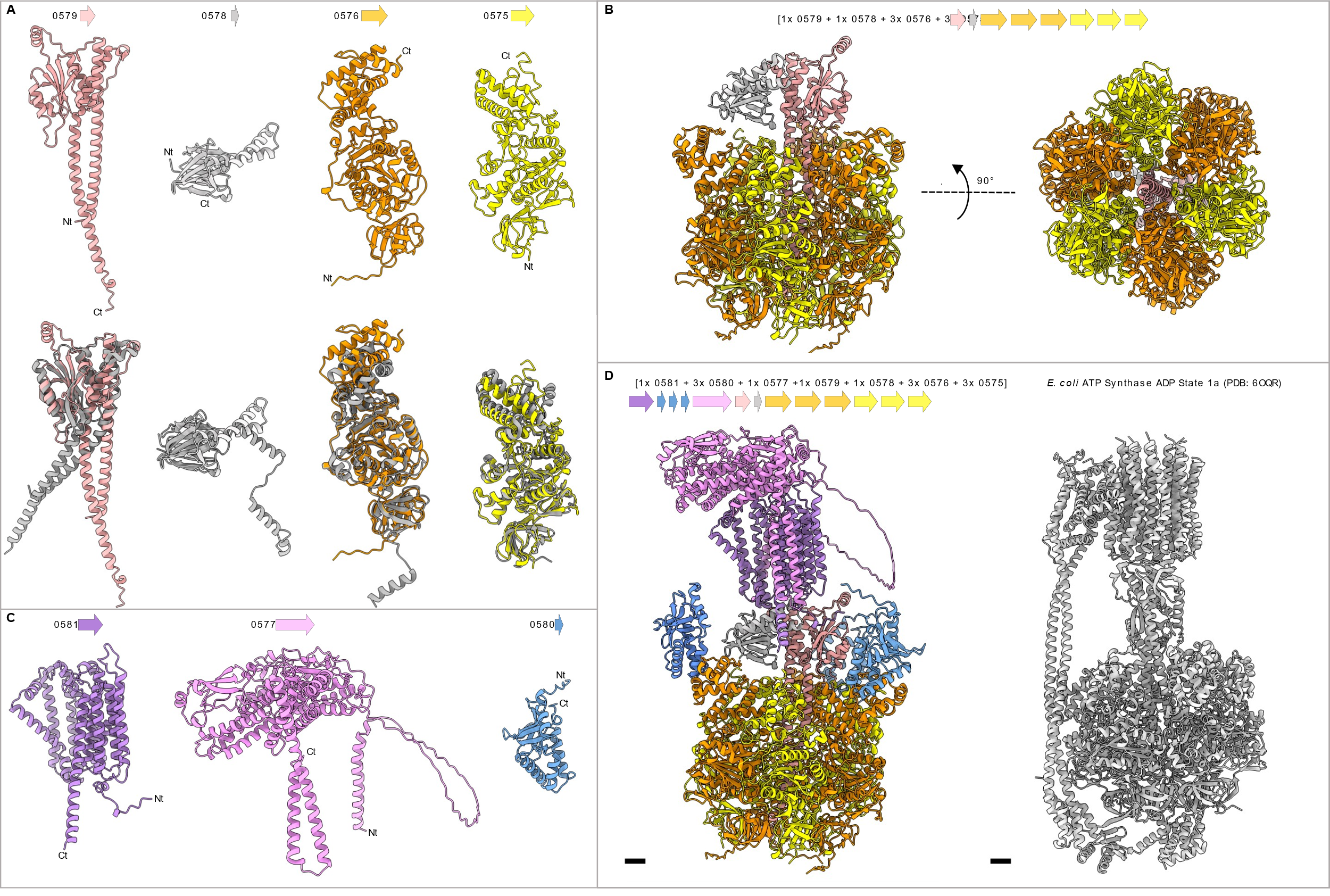

Fig S1

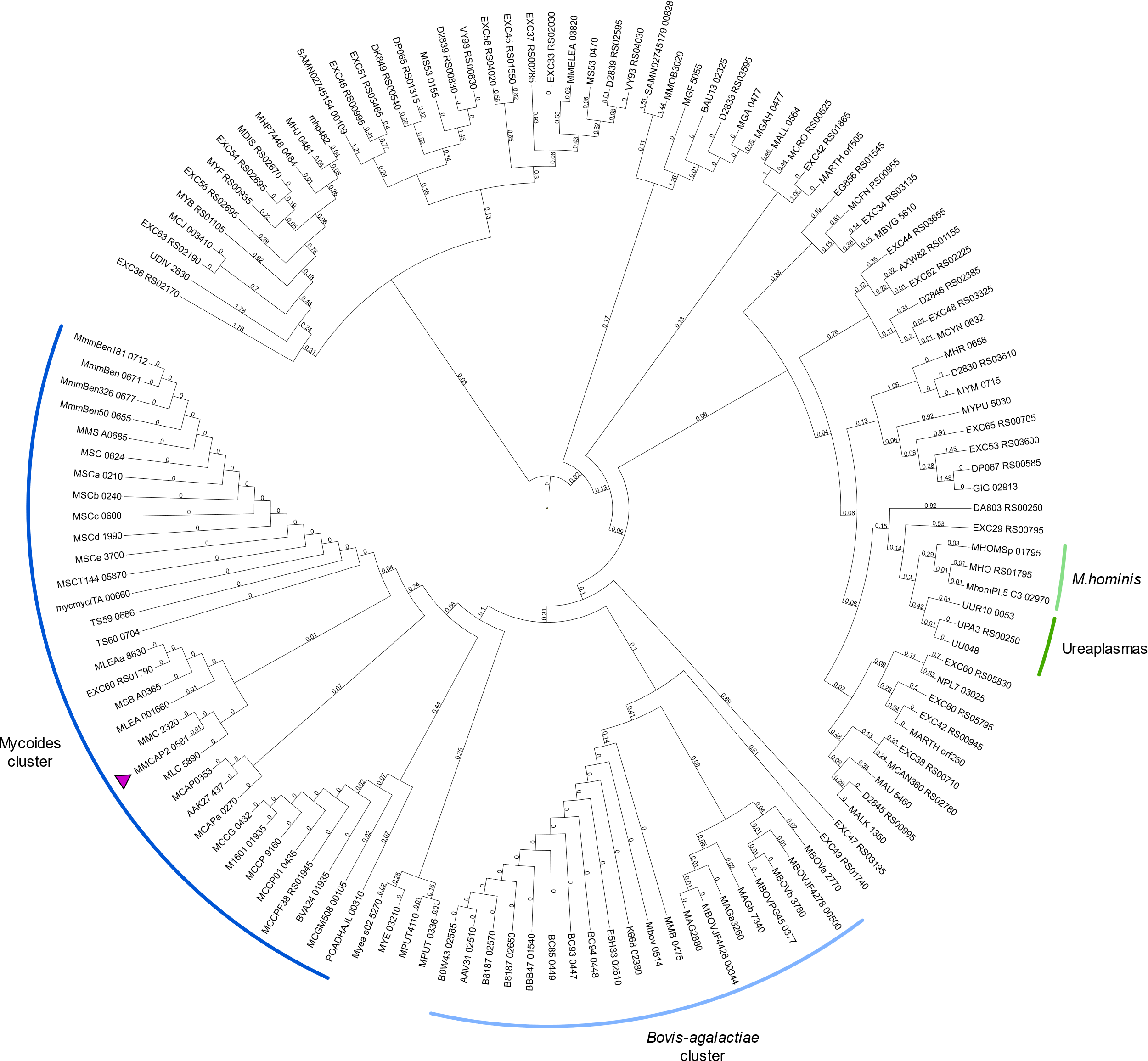

Fig S2

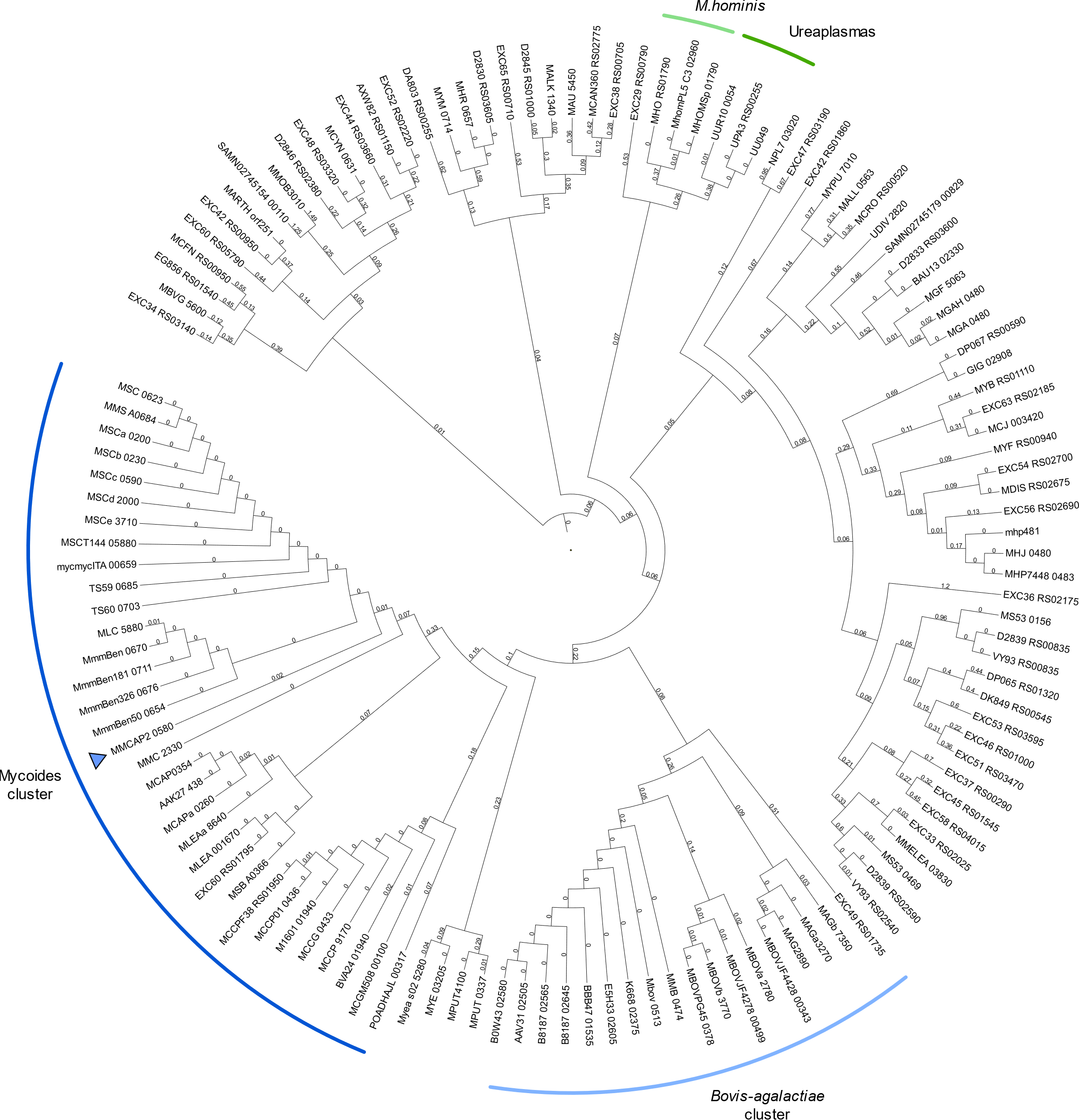

Fig S3

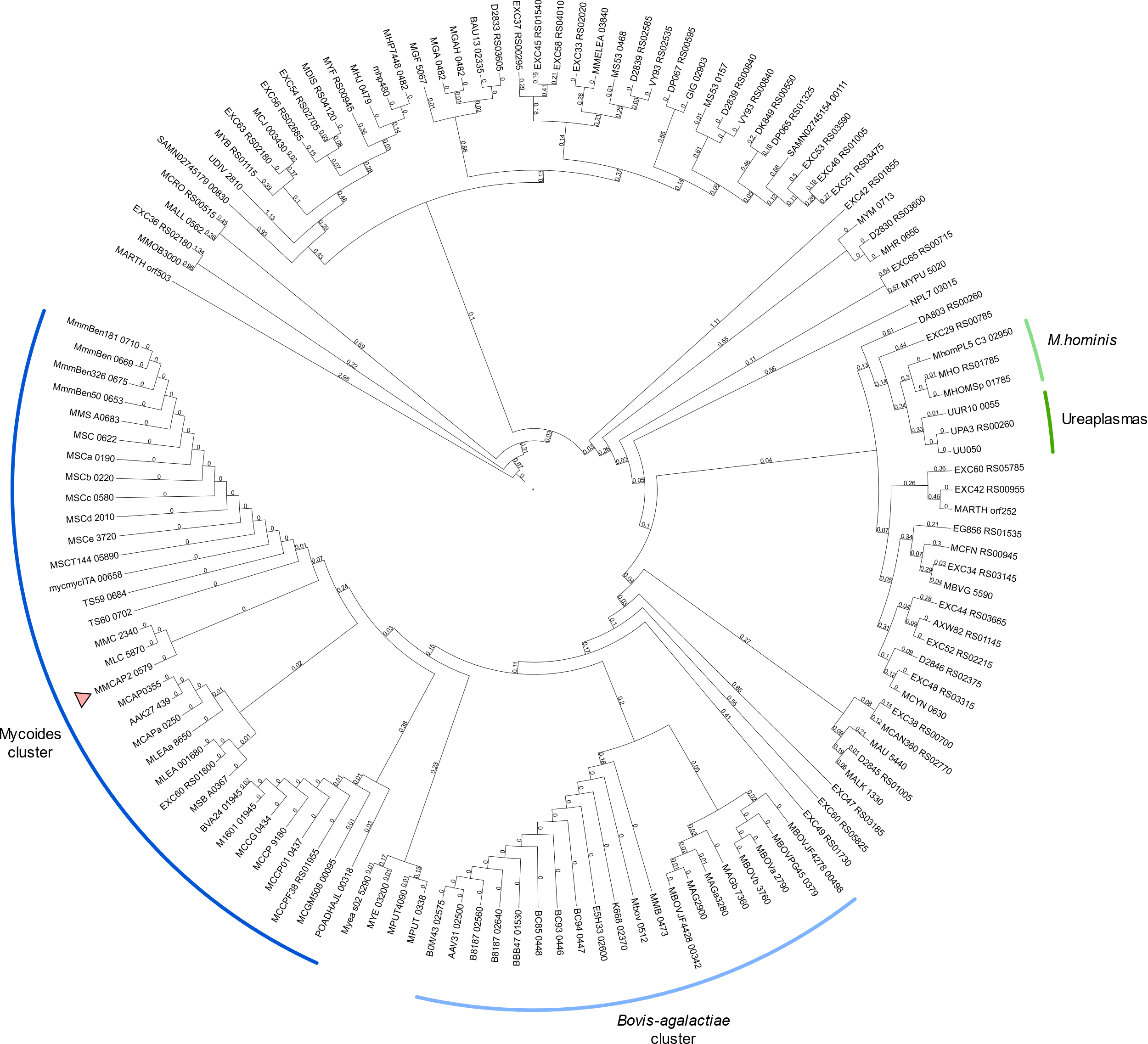

Fig S4

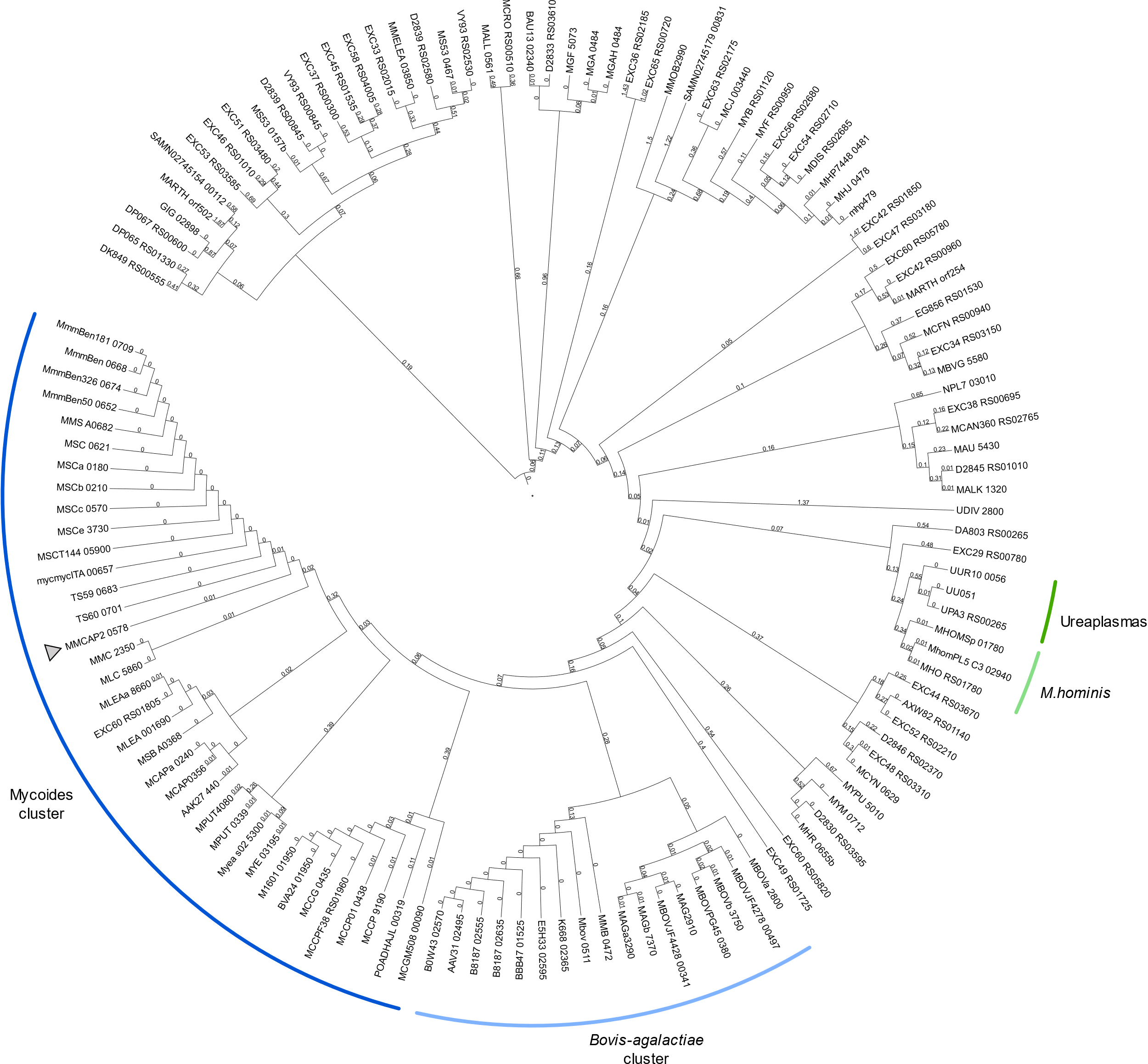

Fig S5

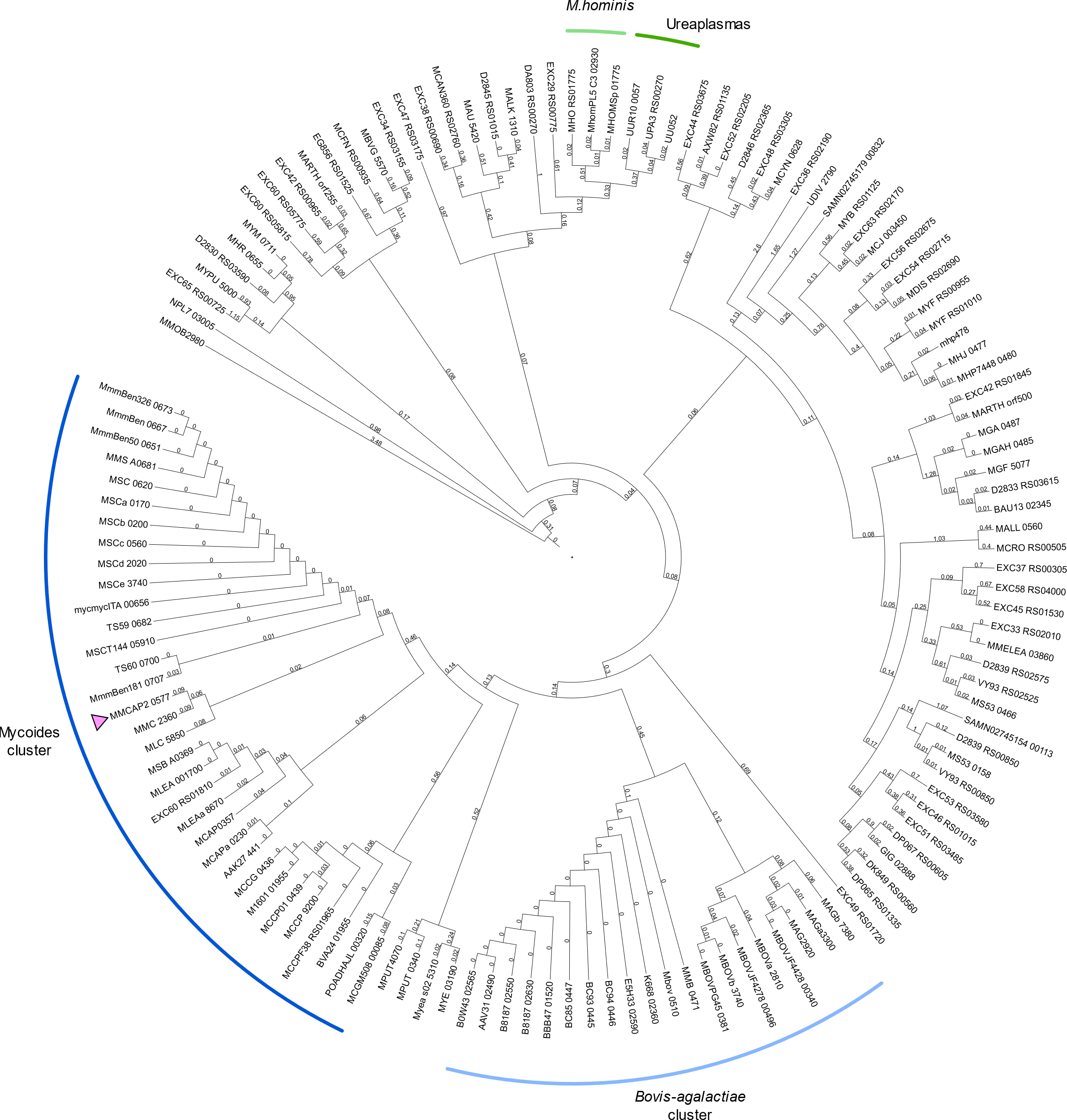

Fig S6

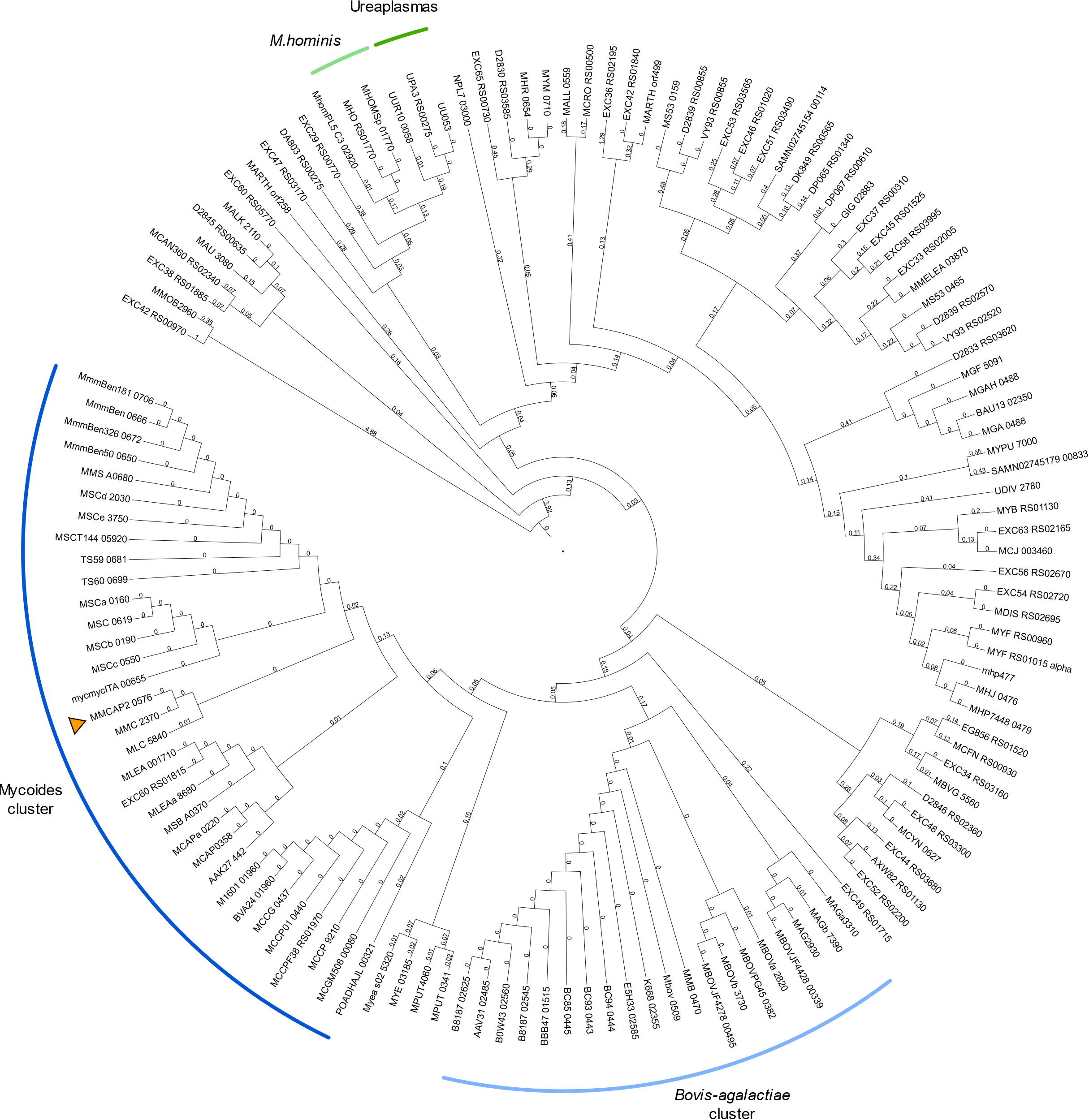

Fig S7

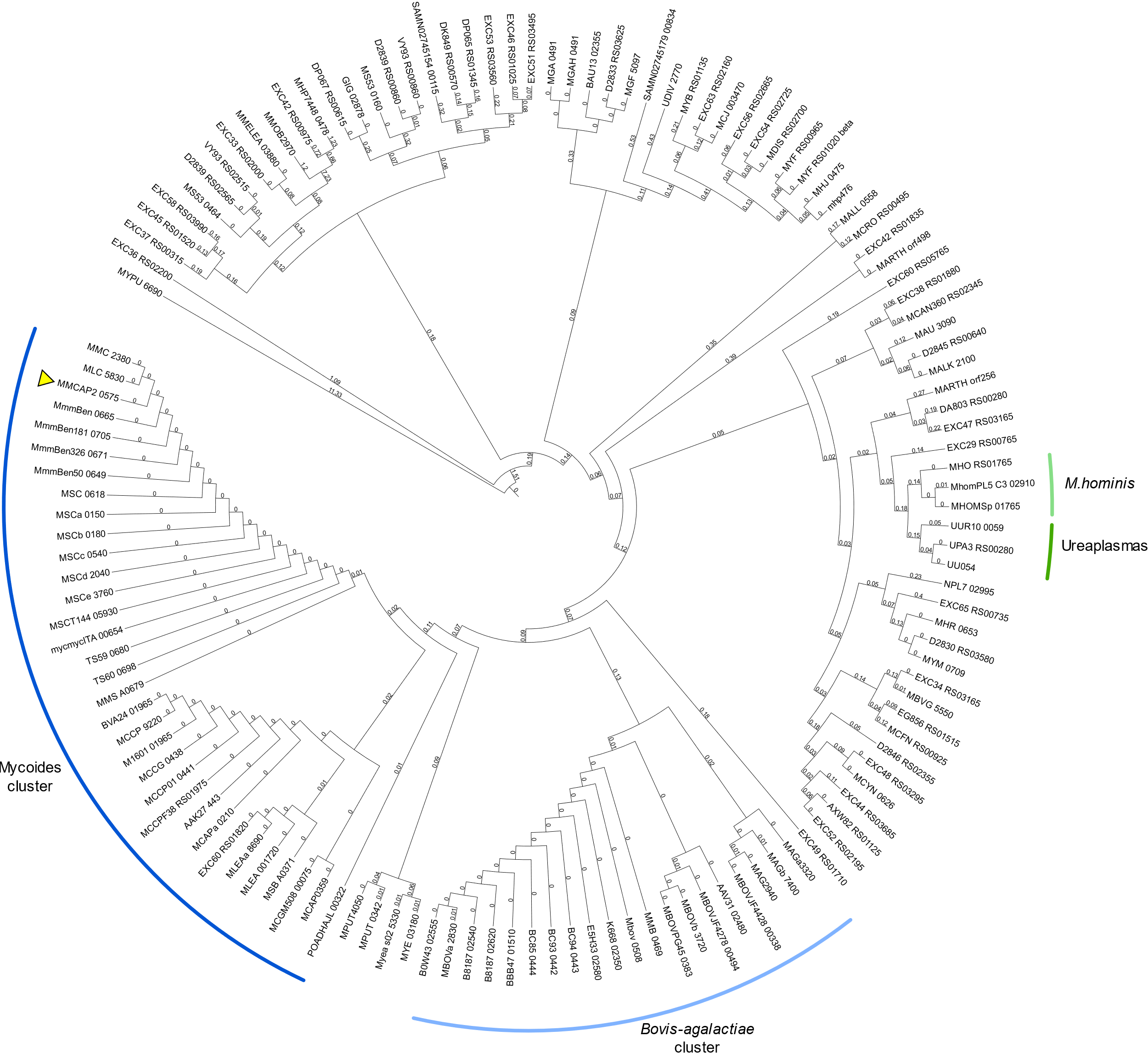

Fig S8

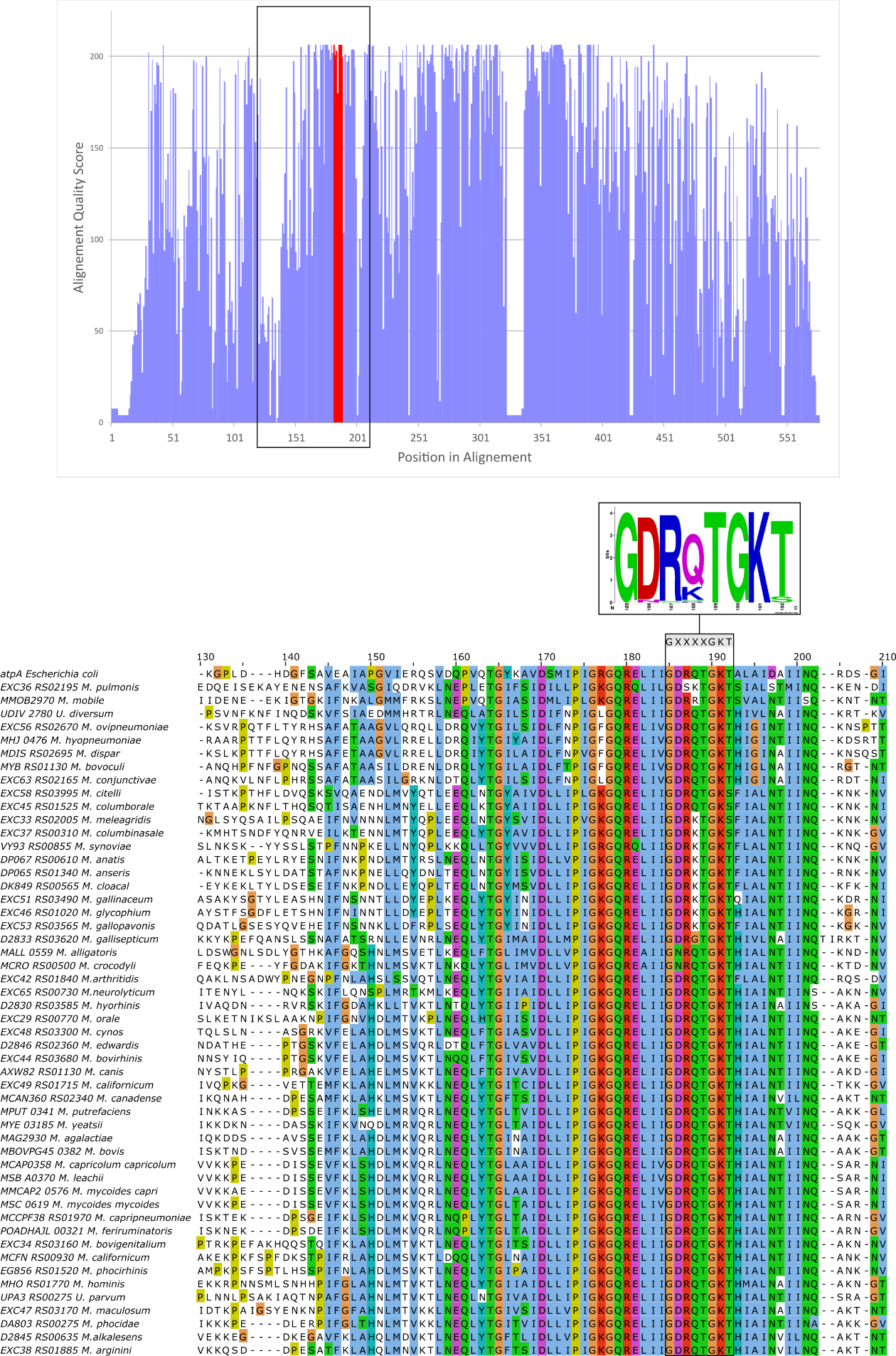

Fig S9

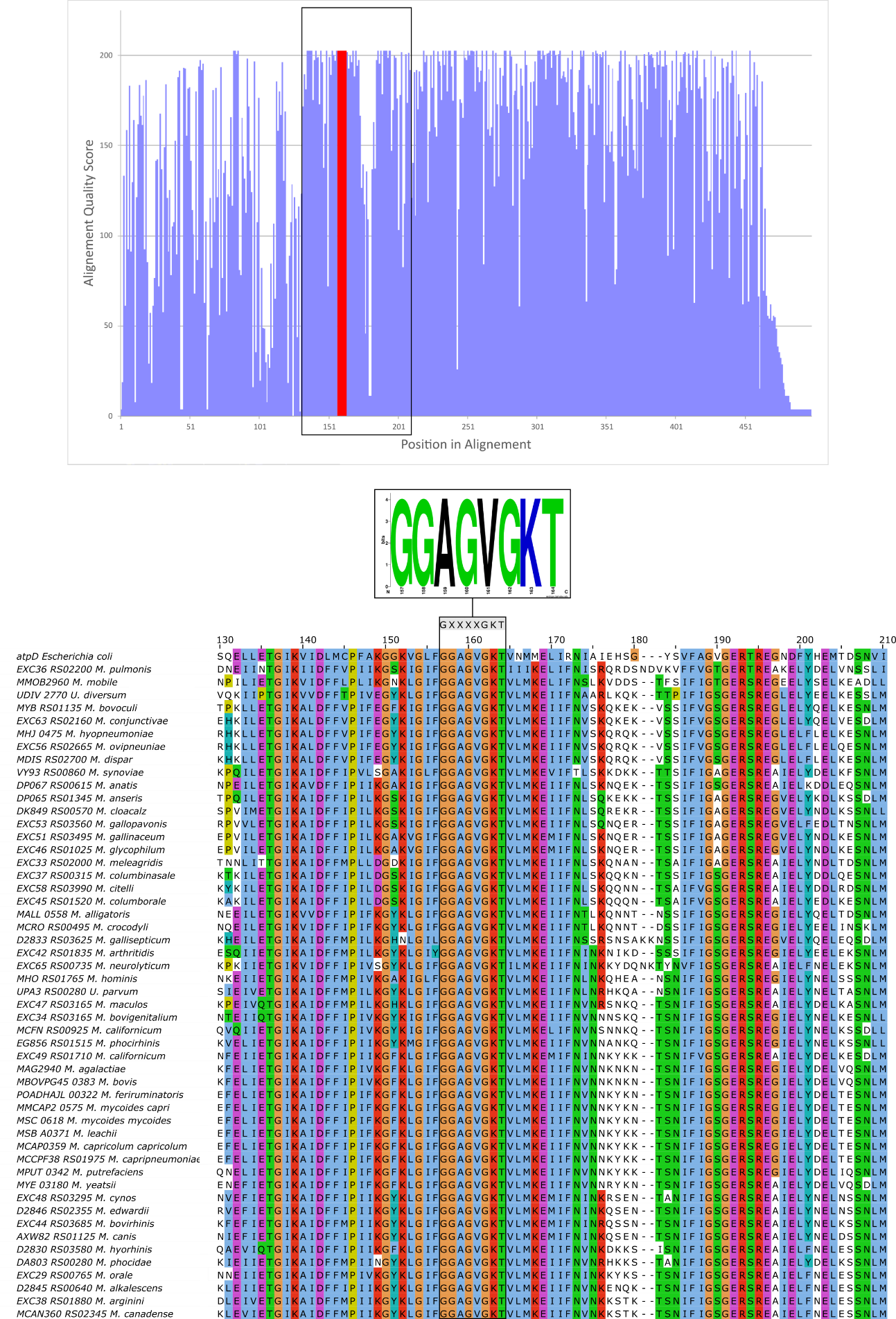

Fig S10

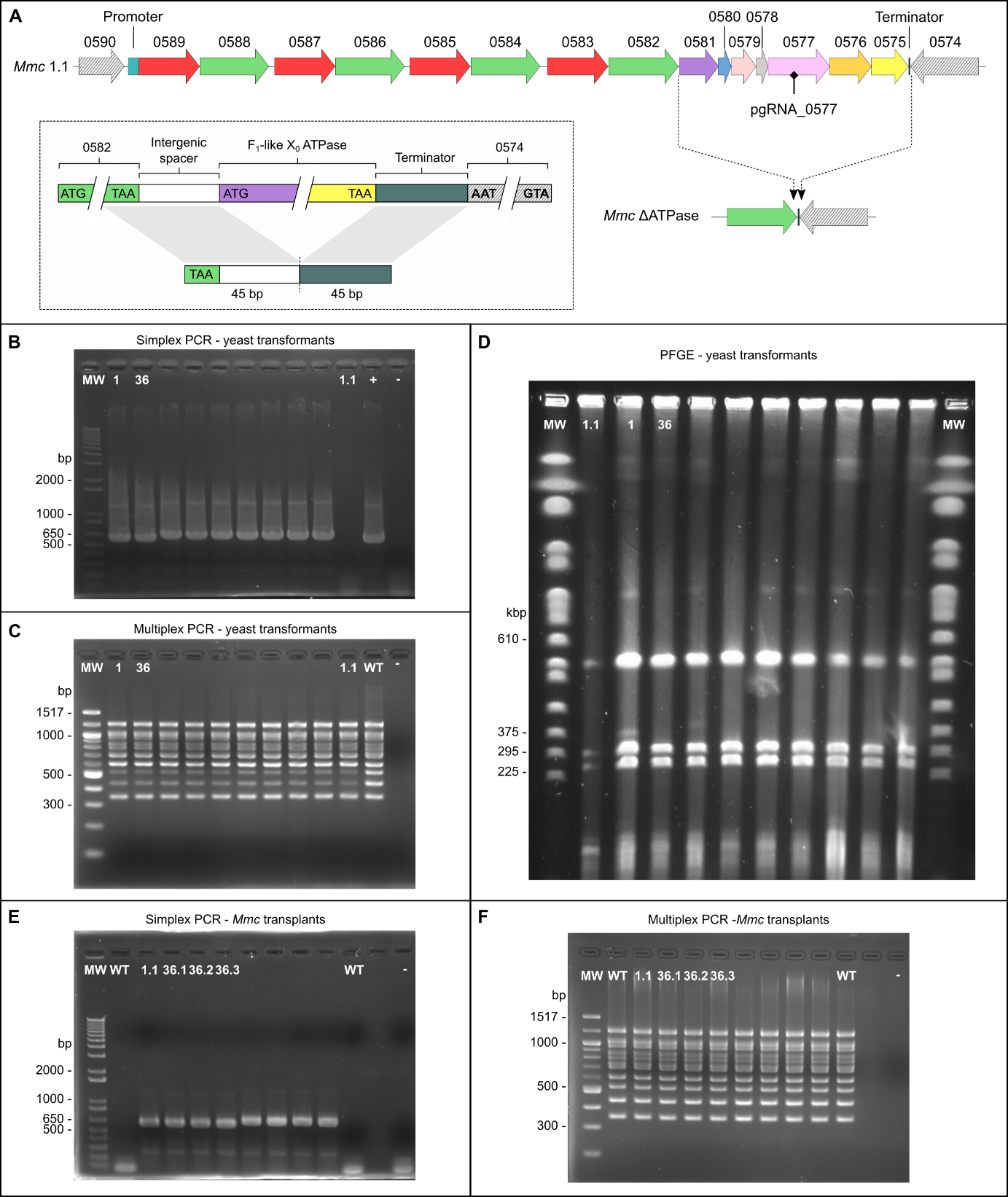

Fig S11

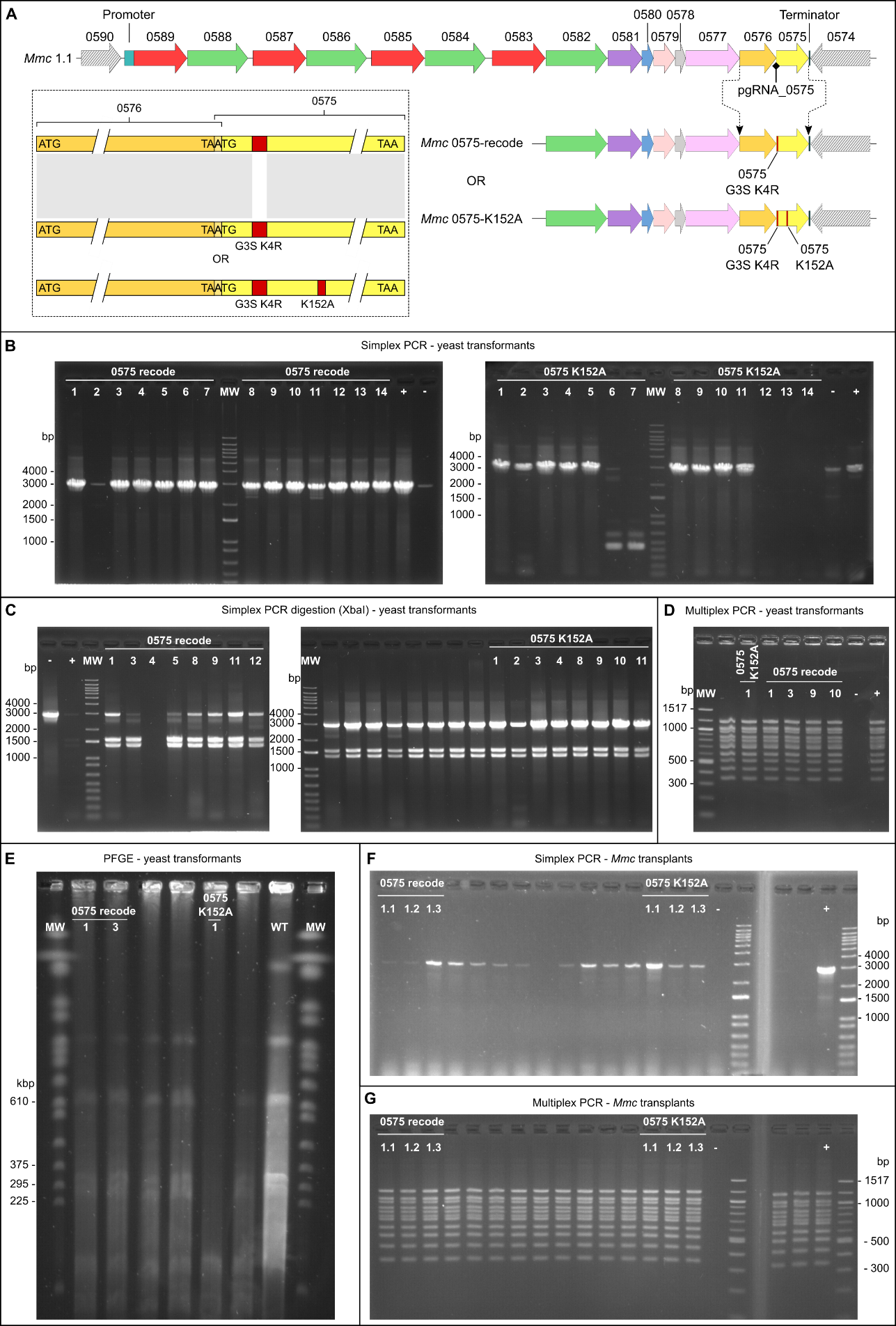

Fig S12

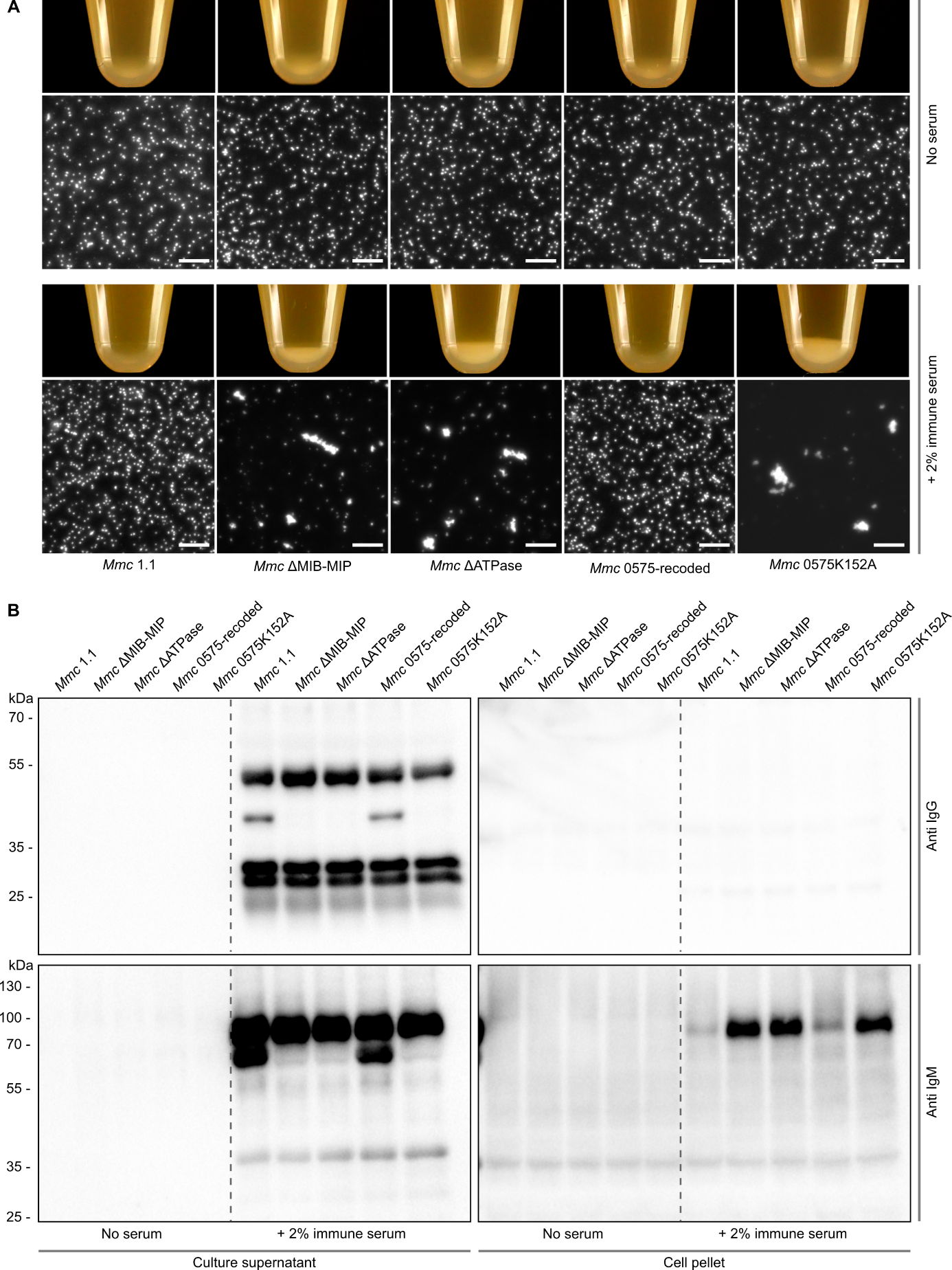

Fig S13

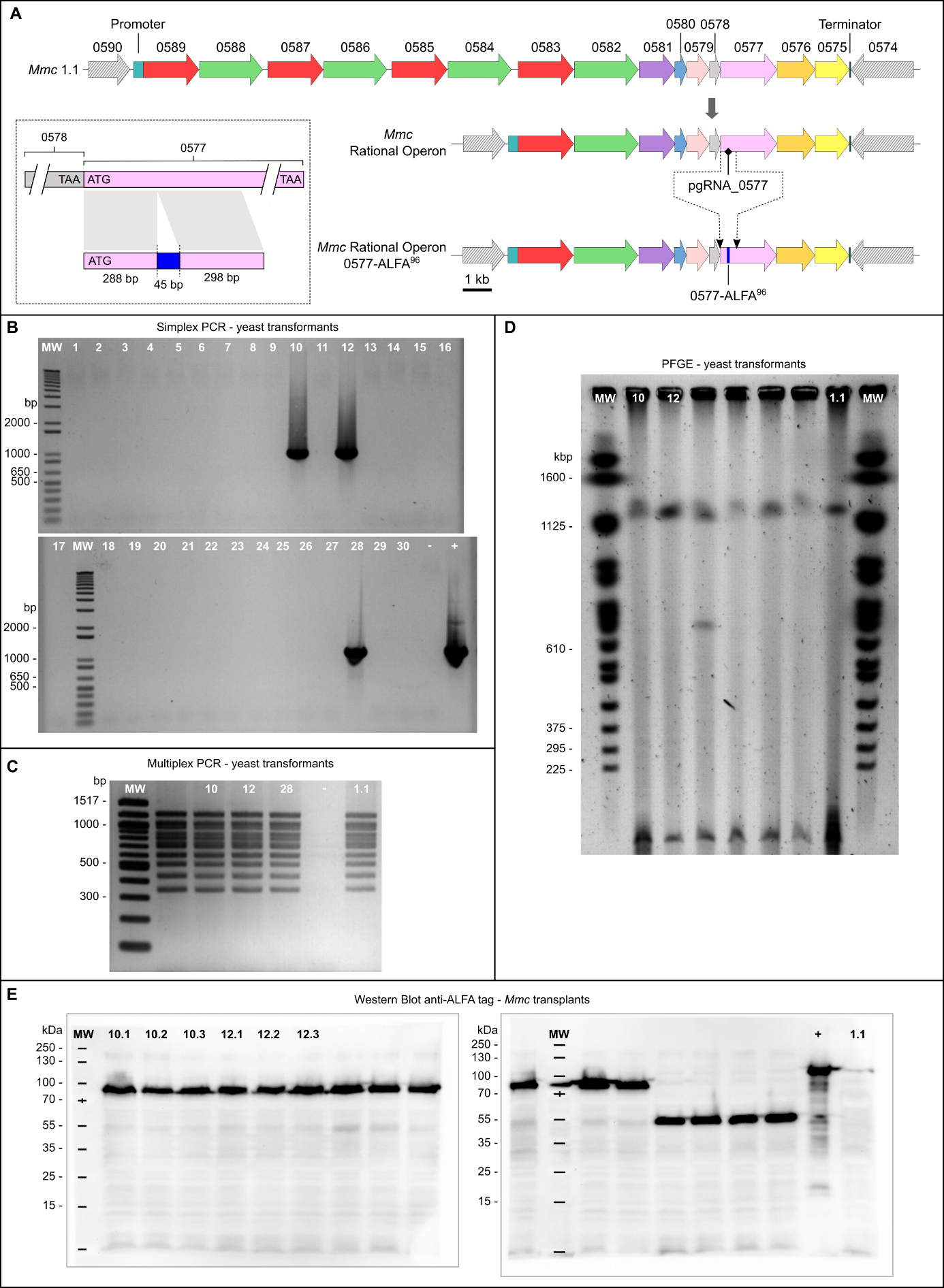

Fig S14

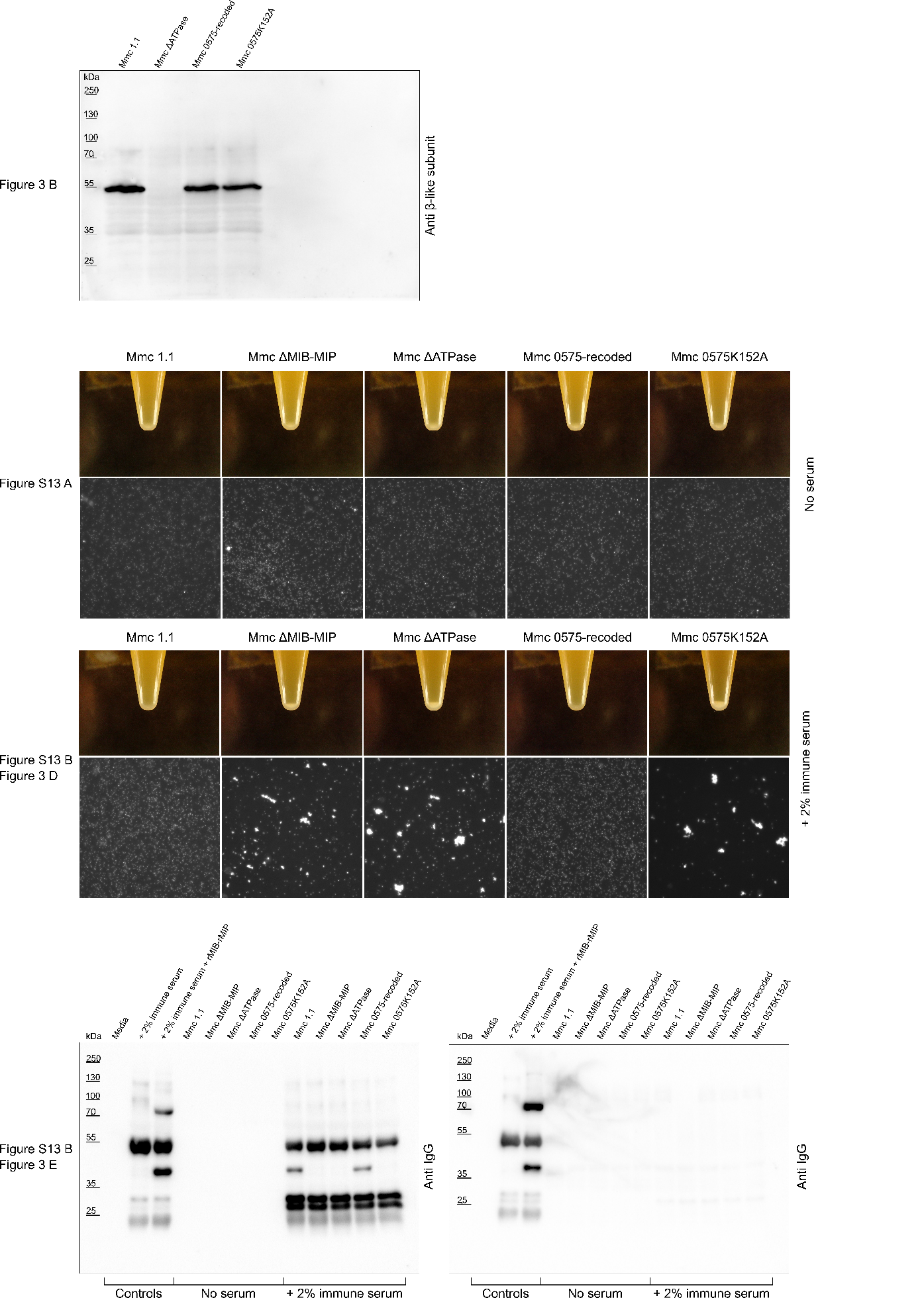

Fig S15-1

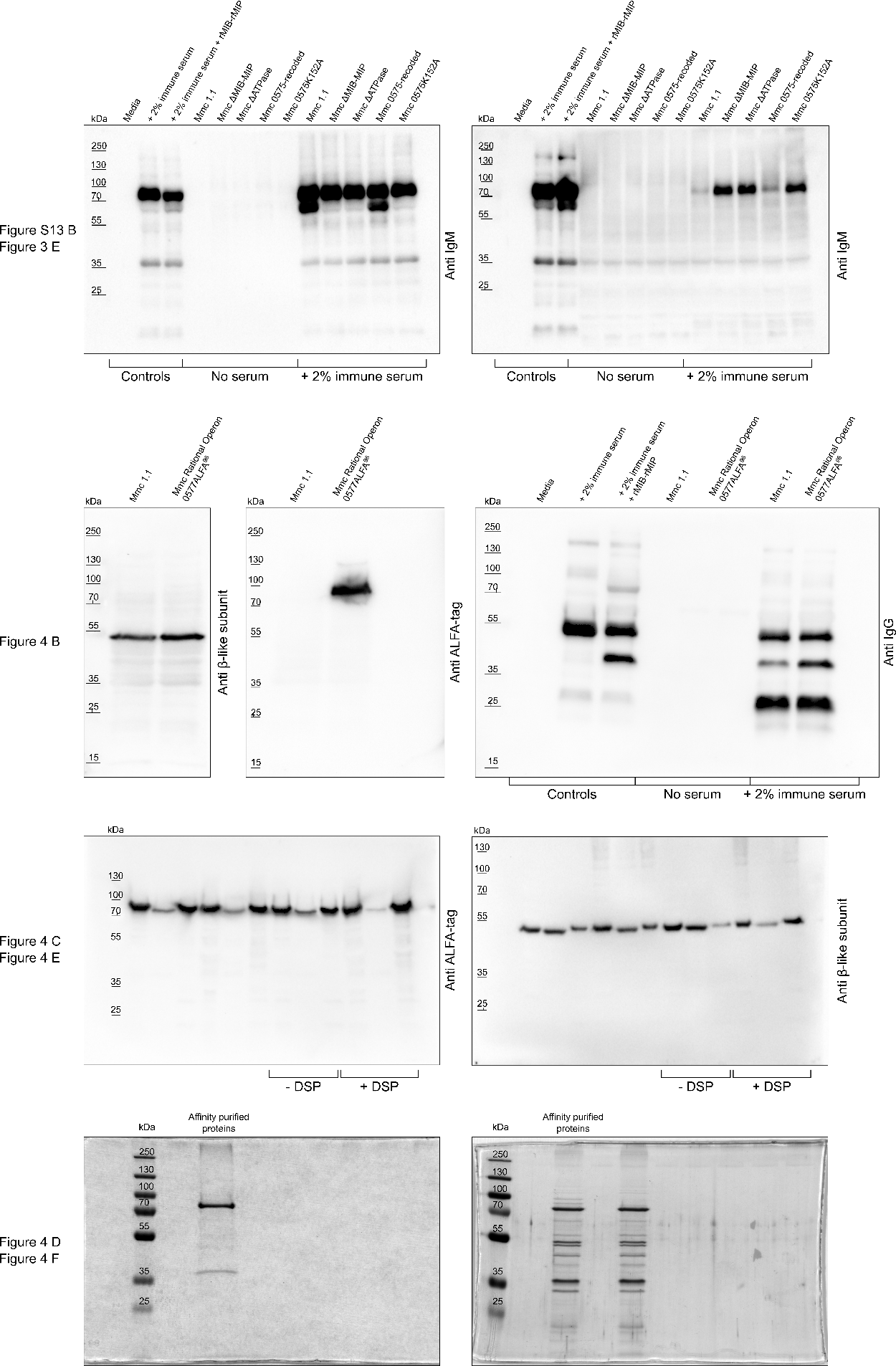

Fig S15-2
